# Metabolic modeling of sex-specific tissue predicts mechanisms of differences in toxicological responses

**DOI:** 10.1101/2023.02.07.527430

**Authors:** Connor J. Moore, Christopher P. Holstege, Jason A. Papin

## Abstract

Male subjects in animal and human studies are disproportionately used for toxicological testing. This discrepancy is evidenced in clinical medicine where females are more likely than males to experience liver-related adverse events in response to xenobiotics. While previous work has shown gene expression differences between the sexes, there is a lack of systems-level approaches to understand the direct clinical impact effect of these differences. Here, we integrate gene expression data with metabolic network models to characterize the impact of transcriptional changes of metabolic genes in the context of sex differences and drug treatment. We used Tasks Inferred from Differential Expression (TIDEs), a reaction-centric approach to analyzing differences in gene expression, to discover that androgen, ether lipid, glucocorticoid, tryptophan, and xenobiotic metabolism have more activity in the male liver, and serotonin, melatonin, pentose, glucuronate, and vitamin A metabolism have more activity in the female liver. When TIDEs is used to compare expression differences in treated and untreated hepatocytes, we see little response in those sex-altered subsystems, and the largest differences are in subsystems related to lipid metabolism. Finally, using sex-specific transcriptomic data, we create individual and averaged male and female liver models and find differences in the import of bile acids and salts. This result suggests that the sexually dimorphic behavior of the liver may be caused by differences in enterohepatic recirculation, and we suggest an investigation into sex-specific microbiome composition as an avenue of further research.

**Author Summary:** Male-bias in clinical testing of drugs has led to a disproportionate number of hepatotoxic events in women. Previous works use gene-by-gene differences in biological sex to explain this discrepancy, but there is little focus on the systematic interactions of these differences. To this end, we use a combination of gene expression data and metabolic modeling to compare metabolic activity between the male and female liver and treated and untreated hepatocytes. We find several subsystems with differential activity in each sex; however, when comparing these subsystems with those pathways altered by hepatotoxic agents, we find little overlap. To explore these differences on a reaction-by-reaction basis, we use the same sex-specific transcriptomic data to contextualize the previously published Human1 human cell metabolic model. In these models we find a difference in flux for the import of bile acids and salts, suggesting a potential difference in enterohepatic circulation. These findings can help guide future drug design, toxicological testing, and sex-specific research to better account for the entire human population.

## Introduction

Male subjects in both animal and human studies are disproportionately used for testing in toxicology studies (Zucker & Beery 2010; Feldman et al., 2019). This discrepancy leads to incorrect assumptions on female drug response as evidenced in the clinic where female patients are more likely than males to experience liver-related adverse events in response to xenobiotics such as acetaminophen, diclofenac, and isoniazid (O’Connor, Dargen, and Jones, 2003; Banks et al. 1995; Ostapowicz et al., 2002). While gene expression differences between males and females have been extensively studied (Yang et al., 2006; Yang et al. 2012; Lopes-Ramos et al., 2020), little is known about how these changes in expression contribute to functional changes that result in this divergent clinical response.

Sexually differential metabolism is key to understanding these responses (Pannala et al., 2019). Transcriptional profiling can provide insight into metabolic genes and how associated functions differ between males and females. A previous study has shown that male and female serum metabolisms have distinct signatures (Mittelstrass et al., 2011), but connecting differences in gene expression and cellular machinery cannot be done on a gene-by-gene basis. To analyze trends across thousands of genes and reactions, a systems-level approach is required.

Genome-scale metabolic models (GEMs) have emerged as a tool for an integrated analysis of the genome, transcriptome, and metabolome. A GEM is a mathematical representation of all known reactions, metabolites, and gene-protein-reaction mappings (GPRs) that characterize intracellular metabolism of a given cell type. The GPRs associate each gene with one or more reactions, so any changes in genome content or in transcription can be evaluated mechanistically with the model. Previous work has investigated sex differences in metabolism (Thiele et al., 2020) and characterized drug-toxicity responses (Rawls et al., 2019) using GEMs, but there is a gap in knowledge in how these factors interact. Transcriptomic data for each case can therefore be used to inform the GEM about which reactions are upregulated or downregulated for a given sex or drug. These differences in gene expression can then be evaluated in the context of GPRs to describe how the flux of each reaction changes in these conditions.

Here, we present an analysis of male and female liver metabolism using tissue-specific and drug-specific gene expression data in the context of a human GEM. Liver-, kidney-, and brain-sourced transcriptomics from the Gene Expression Omnibus (GEO) are used to establish general and liver-specific sex differences in metabolism where the liver acts as the sexually dimorphic tissue of interest, the kidney represents a similarly sexually dimorphic tissue, and the brain functions as a known sexually monomorphic tissue (Yang et al., 2006). We also compare metabolism between hepatocytes with and without exposure to drug using expression data from ToxicoDB (Nair et al., 2020) with particular interest in those metabolic differences found between the male- and female-sourced liver tissue. We then use the male- and female-sourced liver gene expression data to create individual and averaged male- and female-specific GEMs and use flux sampling to illustrate differences in core metabolism and metabolites involved in enterohepatic recycling. Together, these results suggests that sexually dimorphic adverse event frequency may be driven by differences in the gut-liver axis.

## Results

### Women experience liver-related adverse events more frequently than men

We used the Food and Drug Administration’s Adverse Event Reporting System (AERS), a collection of side-effect reports voluntarily sourced from health care professionals and the general public, to quantify sex-specific adverse event frequency (Figure 1). Each report includes information on the event, the sex and age of the patient, other drugs being used, and the time of the incident. We collected reports from this database using AERSMine (Sarangdhar et al., 2016), an application designed to improve the accessibility of AERS. Reports related to liver dysfunction were counted for each quarter from 2004 to 2021 and divided by the total amount of reports for that quarter to account for increasing reporting trends (Figure 1a). We found that not only did more total reports exist for female patients, but that female patients are consistently reported to experience liver-related adverse events more often than their male counterparts (Figure 1b). This result can be explained in part by a previous report of overall increased prescription drug consumption in women (Hales, et al., 2019), so we next investigated sexual-dimorphic response by drug. We compared the ratio of reports for each drug for each sex, only including drugs with greater than 100,000 reports to ensure that the differential effect was robust and not due to small sample size. Unexpectedly, more drugs were disproportionately affecting men, but those that were reported more often in women tended to have a higher fraction of female-specific reports (Figure 1c). It is important to note, however, that not all drugs are used equally by both sexes. For example, alendronic acid is primarily prescribed for osteoporosis and tenofovir disoproxil is used as an HIV treatment, diseases with higher female and male incidence, respectively (Cawthon, 2011; CDC, 2019). Frequently-used pharmaceuticals exhibiting a spectrum of responses indicates that sex is a relevant variable in determining hepatotoxicity, so understanding the sex-based metabolic signature is an important aspect of this problem.

**Figure 1:**
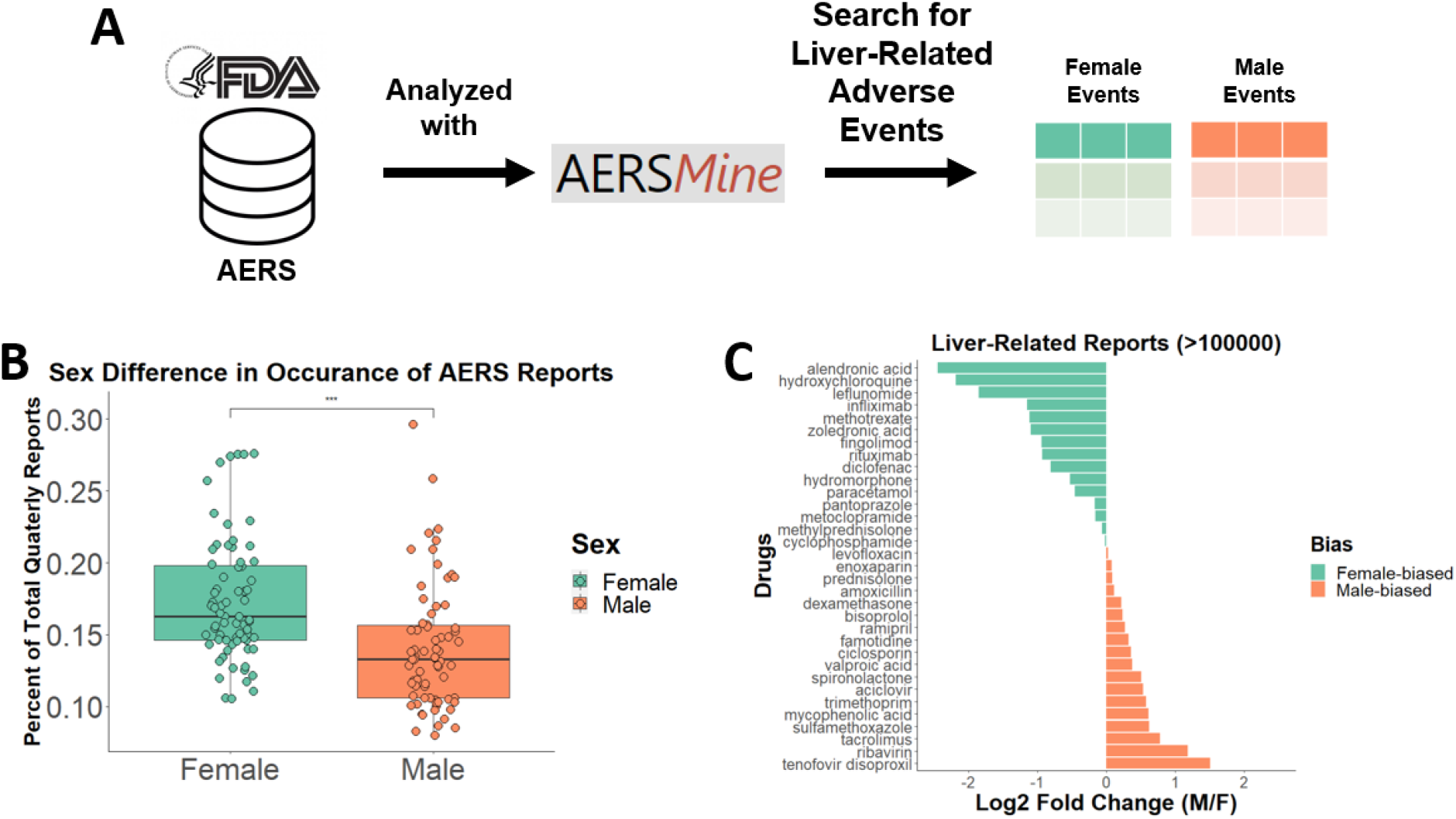
**(A)** The Adverse Event Reporting System (AERS) from the FDA was searched with AERSMine using liver-related terms. **(B)** All reports with sex data were counted for each quarter and compared. *** indicates significance of p-value < .001. **(C)** Reports were then compared for the top reported drugs for all time periods.

### Transcriptome-informed metabolic model indicates a sex- and tissue-specific signature in untreated tissue

Next, we sought to understand differences in functional liver metabolic networks between males and females. To do this, we used microarray data from GEO to characterize gene expression differences between male and female patients not experiencing toxicity. To understand sexual dimorphic metabolism specific to the liver, kidney and brain tissue were also included in this analysis as a comparison. While the kidney is understood to have sex-specific metabolic function, the brain is considered a sexually monomorphic tissue with respect to gene expression (Yang et al., 2006).

For each tissue type, differentially expressed genes (DEGs) between the sexes were calculated (FDR < 0.1) (Figure 2a). We then used the publicly available GEM, Human1 (Robinson et al., 2020) to provide context for the functional impact of the differentially expressed genes. Human1 accounts for the function of over 13000 reactions, 8300 metabolites, and 3600 genes with GPRs linking these genes and reactions. Using a GEM in conjunction with microarray data allows us to provide a reaction-centric analysis with Tasks Inferred from Differential Expression (TIDEs) (Dougherty et al. 2021) instead of the traditional gene-by-gene view (Figure 2c). Using the subsystems defined by Human1, we can assign weights to each reaction in the subsystem dependent on the log fold change from the DEGs and the GPRs from the GEM to generate a task score for that subsystem. This task score can then be compared to randomized weights for each reaction to determine if the task score is significantly higher or lower (male- or female-biased) for that subsystem.

**Figure 2:**
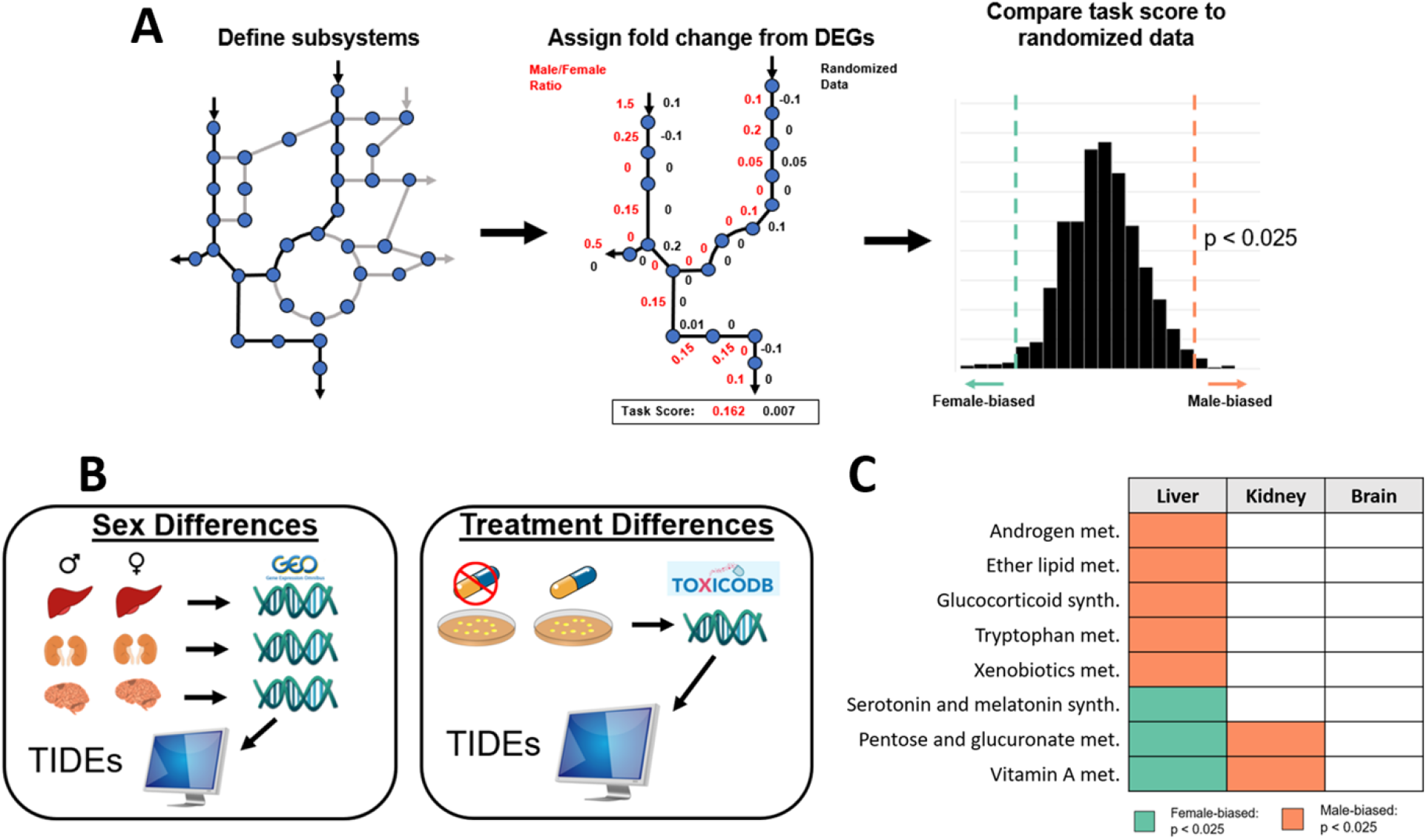
**(A)** Tasks Inferred from Differential Expression (TIDEs) is summarized. First, reactions are selected to be part of a metabolic subsystem. Expression data and GPRs are then used to assign log2 fold changes as weights to each reaction. The task score is then calculated as the average weight of the subsystem and compared to randomized fold changes from other genes in the data to determine significance. **(B)** This method is used to compare sex-specific untreated liver, kidney, and brain expression data as well as treated and untreated cultured hepatocyte expression data. **(C)** 8 subsystems are found to be differentially expressed between the male and female samples in a liver-specific manner while the brain shows no dimorphic behavior.

Of the 135 subsystems defined by the GEM, 11 in the liver tissue, 23 in the kidney, and 0 in the brain were found to be significantly different between the male and female patients. The gene expression data for the male and female brain tissue did not have differences in TIDEs, suggesting highly similar metabolic function of the sexes in this tissue type. The expression profiles for the liver and kidney tissue, however, did have significant sex-biased TIDEs. Of the sex-biased tasks, only two were found to be upregulated by the same sex between the liver and kidney: Acylglyceride Metabolism and Steroid Metabolism pathways were upregulated in males. Previous literature is consistent with increased very low-density lipoprotein and triglyceride production in male livers (Palmisano et al., 2018) and a higher rate of steroid metabolite secretion in men (Raven & Taylor, 1996). Of the other 8 subsystems significantly different in the liver, 6 are only significant in the liver and 2 are biased in opposite sexes (Figure 3a). Vitamin A storage in rats has been shown to be higher in female livers and male kidneys (Booth, 1952), agreeing with these results. Additionally, androgen receptors and glucocorticoid receptors have been shown to be correlated in mouse livers (Kroon, Pereira, & Meijer, 2020), and rat liver tissue has previously exhibited an increase in tryptophan 2,3 dioxygenase activity in response to glucocorticoids (Danesch et al., 1983), explaining how these three metabolic subsystems would all be upregulated in male liver tissue. Tryptophan also acts as a precursor to serotonin and melatonin. With more of the amino acid being used by male tryptophan 2,3 dioxygenase, an enzyme which produces no products involved in serotonin and melatonin metabolism, less tryptophan is available for serotonin and melatonin biosynthesis, explaining the relative increase in female activity for this task. Previous research has also shown that ether lipid levels in female serum tend to be higher than in male serum (Lim et al., 2020), suggesting a possible increase in ether lipid metabolism in men. Pentose and glucoronate interconversions are more active in the liver in females and in the kidney for males. It is interesting to note that part of this subsystem includes metabolites that are products of UDP-glucuronosyltransferases (UGTs), enzymes that encapsulate part of phase II detoxification (Shipkova et al., 2003), indicating a potential difference in how toxic metabolites are handled.

**Figure 3:**
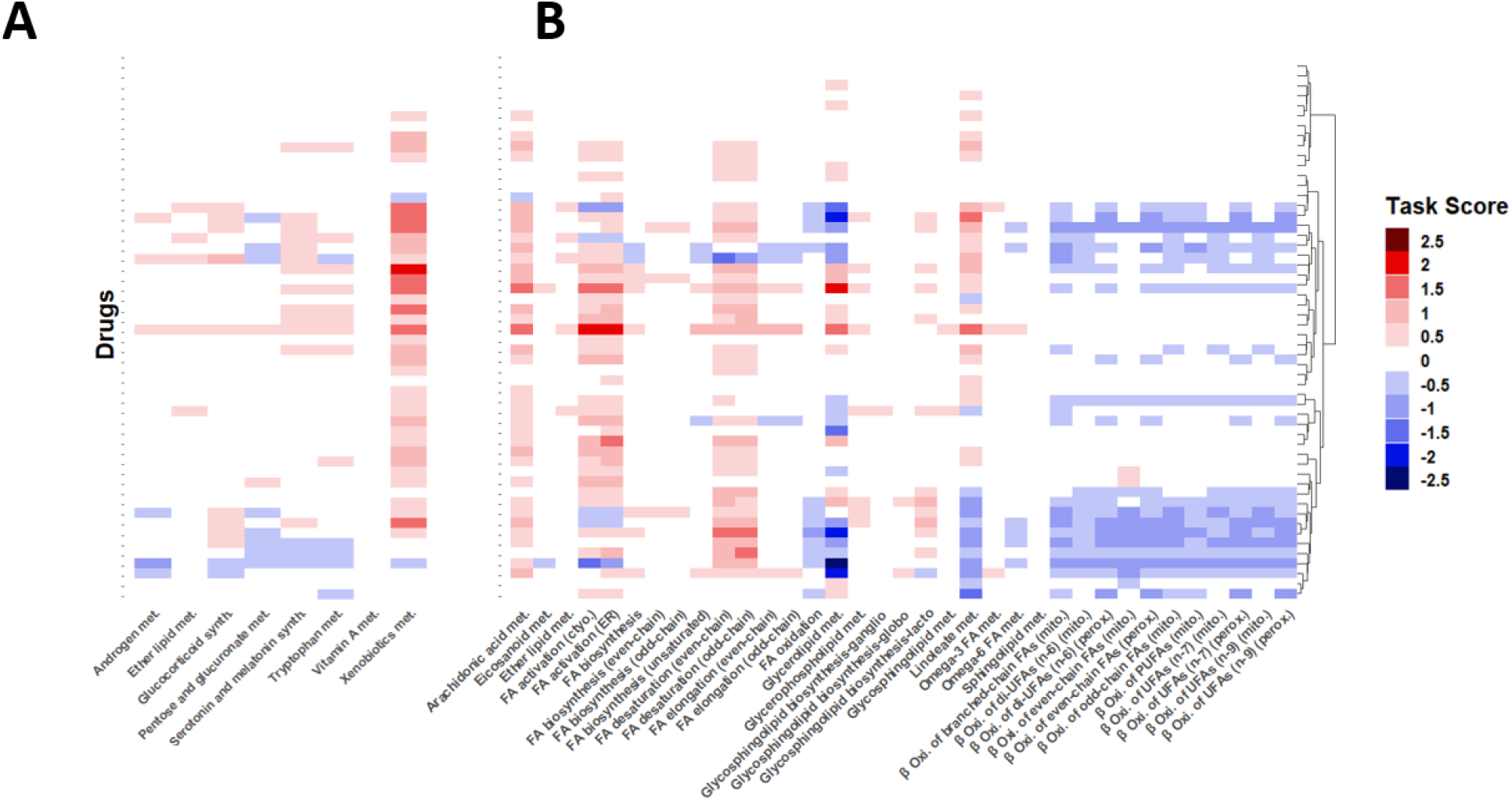
The results of a task score comparison of non-treated hepatocytes and drug-treated hepatocytes with TIDEs are shown with each row representing a different treatment and each column representing a different subsystem. **(A)** Here we show the tasks shown to be sexually dimorphic in the previous analysis. In this comparison, only Xenobiotic Metabolism appears to have a significant response. **(B)** Here we look at subsystems related to lipid metabolism. These subsystems show altered behavior when treated with hepatotoxic agents especially when compared to those subsystems compared in (A).

### Hepatocyte expression data suggests the metabolic response to hepatotoxic pharmaceuticals does not overlap with sex-biased metabolism

Using gene expression data from ToxicoDB (Nair et al., 2020), we used TIDEs to provide context to the DEGs between the treatment and no treatment conditions. We clustered drugs by their task score, a measure of the significance of differences in metabolic tasks (see Methods), and find that drugs group mainly by their effect on lipid metabolism (Supplementary Figure 1). There is no clear clustering by indication, toxicity, or sexually dimorphic nature, and in fact, when we look at just the subsystems found to be significant between healthy male and female livers, xenobiotic metabolism is the only task with a significant effect (Figure 3a). Most drugs appear to increase the expression of genes related to xenobiotic metabolism, while the other sexually differential subsystems have no strong bias in either direction. Many of the subsystems in lipid metabolism, however, have a unidirectional effect (Figure 3B). Arachidonic acid metabolism, fatty acid activation, and fatty acid desaturation tend to increase in activity while beta oxidation of fatty acids tends to decrease in activity. Glycerolipid metabolism and linoleate metabolism force many of these drugs into clusters depending on whether the drugs cause an increase or decrease in expression of these subsystems. Increased lipid storage has been previously reported to be a result of drug-induced toxicity (Begriche et al., 2011), supporting our results of a decrease in beta oxidation of fatty acids. While many drugs interact with lipid-related subsystems, some drugs have a limited effect on any subsystem, indicating that hepatotoxic side effects of all drugs cannot be described with gene expression data alone.

### Flux sampling of individual sex-specific models

While a subsystem-level view can help explain broad differences in metabolism, a more refined comparison of metabolism may provide additional insight. To understand how male and female liver metabolism differ on a reaction-by-reaction basis, we used the same liver-specific gene expression data from TIDEs with Reaction Inclusion by Parsimony and Transcript Distribution, or RIPTiDe (Jenior et al., 2020). RIPTiDe utilizes gene abundances and overall flux parsimony to prune reactions that do not reflect the cell’s transcriptome in a given context, in this case male or female liver (Figure 4a). With this method, we used each individual transcriptome to create sample-specific models. We tested various objective fractions and removed those models which could not achieve at least 40% of the original maximum flux, leaving 586 female and 218 male models to analyze. Each model was sampled 110 times to reach the required sampling depth required for RIPTiDe.

**Figure 4:**
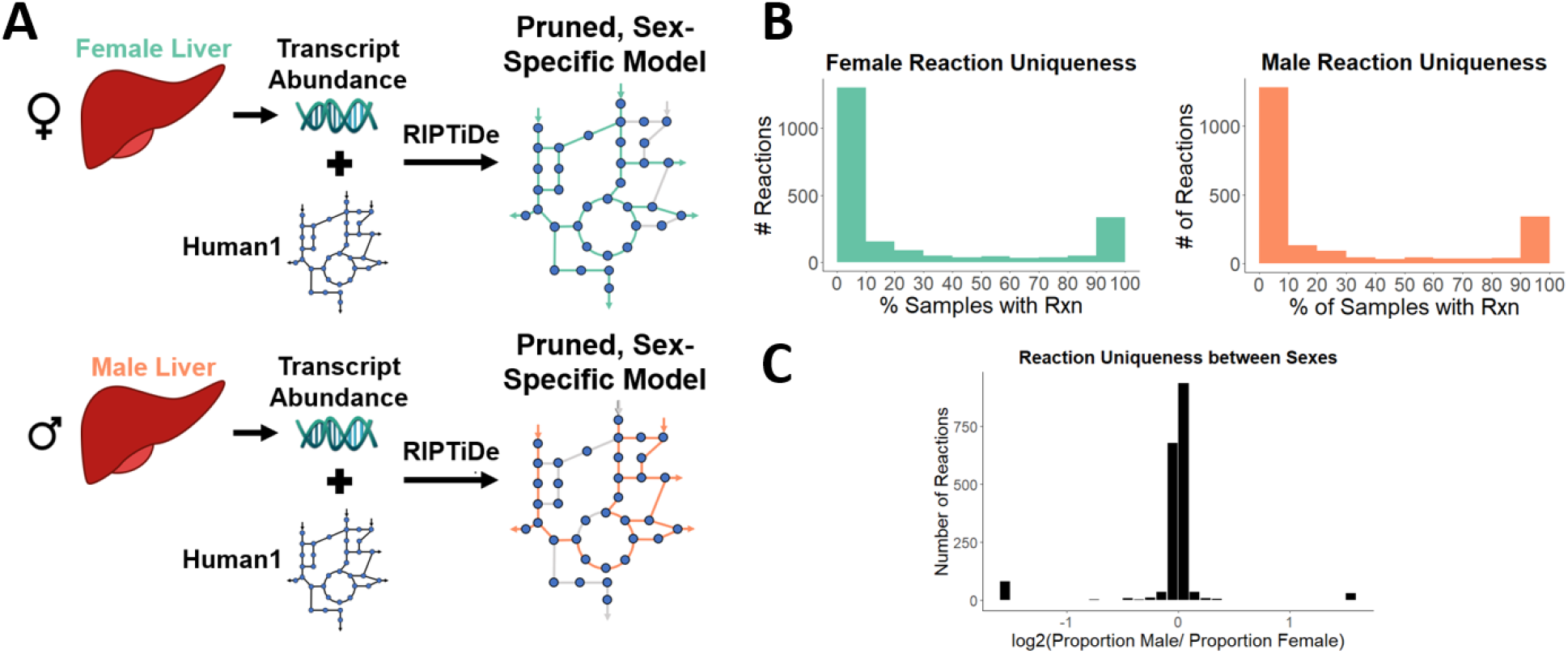
**(A)** RIPTiDe can be used to create sex-specific models from transcriptomic data and an existing model by mapping gene abundance to reactions and pruning those that lack evidence of expression. **(B)** Reactions are checked for uniqueness within each sex. The number of sex-specific samples with a specific reaction is divided by the total number of samples for that sex, and that reaction is placed in the appropriate bin. This process is repeated for every reaction in each set of samples. **(C)** The proportion of male samples that contain a specific reaction was compared to the corresponding proportion contained in female samples to determine uniqueness between sexes. Bars at either extreme represent reactions specific to each sex.

When looking at variability within each sex, we see that reactions are frequently either a part of the core metabolism or are present in very few of the models (Figure 4b). A similar distribution can also be seen when comparing between sexes, where most reactions are found in near-equal proportions in each sex with the second largest group being sex-specific reactions (Figure 4c). Specifically, we found that the male and female models contained 51 and 246 unique reactions, respectively, that are in at least 10% of the samples for each sex. These unique reactions tended to group into specific pathways: purine metabolism in females (Table 1) and transport reactions for males (Table 2). These groups of reactions suggest that while genes directly controlling these subsystems do not result in significant differences in tasks, there are systemic differences in each sex that contribute to differential metabolism.

**Table 1:** Top Unique Subsystems based on Number of Unique Reactions in Female Models.

| Subsystem | # of Unique Female Reactions |
| --- | --- |
| Purine metabolism | 28 |
| Pyrimidine metabolism | 19 |
| Glycolysis / Gluconeogenesis | 12 |
| Nucleotide metabolism | 10 |
| Pentose phosphate pathway | 5 |

**Table 2:** Top Unique Subsystems based on Number of Unique Reactions in Male Models.

| Subsystem | # of Unique Male Reactions |
| --- | --- |
| Transport reactions | 21 |
| Carnitine shuttle (mitochondrial) | 2 |
| Fatty acid oxidation | 2 |
| Nucleotide metabolism | 2 |
| Exchange/demand reactions | 1 |

### Averaged sex-specific models suggest differences in enterohepatic recycling which may impact the hepatotoxic effects of drugs

To characterize broad differences between sexes, we used RIPTiDe with averaged gene expression to create a male and female metabolic model. Since previous literature has suggested that enterohepatic recycling is a relevant factor in hepatotoxicity (Talevi & Bellera, 2022), we decided to compare the bile acids in our models as they provide a comparable set of metabolites to analyze differences in this recycling. The liver synthesizes primary bile acids and conjugates them into bile salts (Cheng, Mah, & Seluakumaran, 2021) which are then transported into the small intestine (Figure 5a). The conjugated bile salts can then be transported into the portal vein, or deconjugated back into a bile acid by the gut microbiome. The resulting primary bile acid could then be transported into the portal vein and back to the liver, where the cycle begins again. Many drugs follow a similar process following glucuronidation, leading to longer half-lives than would be expected. In this analysis, we focus on cholate and its conjugate, glycocholate. We find that the male liver GEM imports significantly more cholate (Figure 5b) while the female liver GEM relies more on glycocholate (Figure 5c). This result suggests that female livers are metabolizing conjugated bile salts at a higher rate than deconjugated bile acids, pointing to a difference in how the acids are deconjugated, how they are exported into the blood, or how they are imported into the liver. A similar mechanism could act on drugs and metabolites as well, leading to larger and more toxic concentrations of injury-inducing agents.

**Figure 5:**
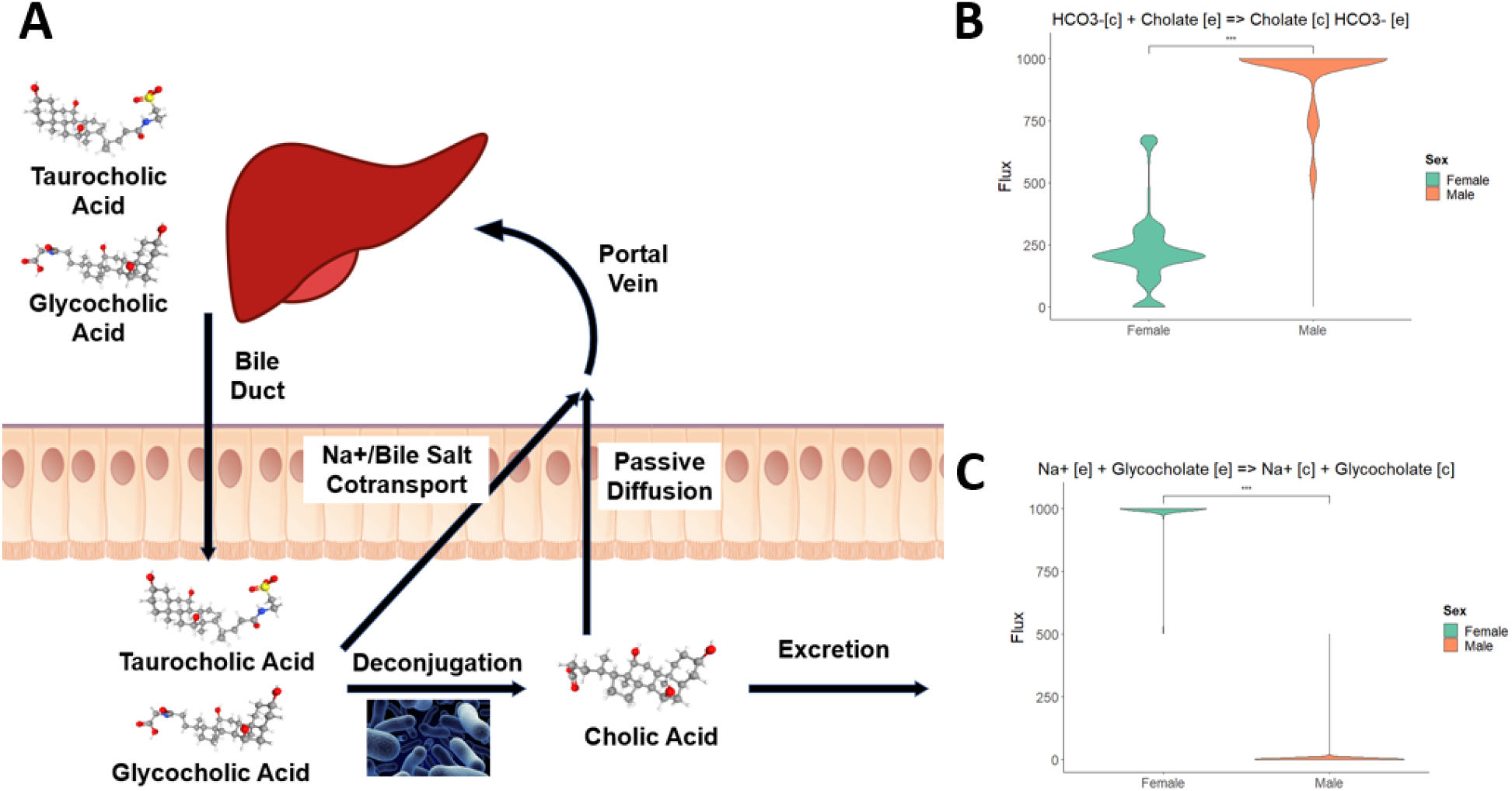
**(A)** Enterohepatic recycling begins with the excretion of conjugated metabolites (here bile acids) into the bile. These metabolites then enter the gut to aid in digestion and can be either reabsorbed by the gut, deconjugated by the gut microbiome, or excreted. Once conjugated, the new metabolite can still be reabsorbed or excreted. Once reabsorbed, the recycled agents can enter the portal vein to once again be taken up by the liver. **(B)** Sex-specific models show a preference of deconjugated bile acids in males and **(C)** conjugated bile acids in females.

## Discussion

We present an analysis of the current landscape of female-biased adverse event reporting, the sexually dimorphic profile of liver metabolism, the hepatocellular response to toxic pharmaceuticals, and differences in sex-specific GEMs. We find that sex as a biological variable is critical to describe the flow of metabolites through the hepatocyte and that there is little overlap between those subsystems which exhibit sex differences and those which experience changes in response to drug. Of the 135 subsystems defined in the Human1 metabolic network reconstruction, 11 in the liver, 23 in the kidney, and 0 in the brain were found to differ based on sex, suggesting that the kidney may have more sex-specific differences in functional metabolism than other tissue types. This result suggests a different conclusion compared to previous literature in mice (Yang et al., 2006) though this result may be an artifact of the available data for this investigation. In the human liver dataset we used for our analysis, the participants are all classified as obese and therefore may be experiencing liver-related health issues that dilute the effect of sex, leading to potentially conflicting results.

In the liver, we find that xenobiotic metabolism was more active in untreated males, while pentose and glucoronate interconversions were female-biased. This result points to a difference in pretreatment gene expression, which may result in different initial responses of phase I and phase II metabolism to hepatotoxic drugs. There is disagreement in previous reports about which enzymes and subsystems are upregulated in each sex (Chu, 2014; Franconi & Campesi, 2014; Waxman & Holloway, 2009), and an up-to-date, focused investigation of these properties could enable the prediction of sex-specific hepatotoxicity as well as suggest sex-specific therapeutic dosing in clinical practice.

Our analysis into the response of hepatocytes to cytotoxic drugs showed a general decrease in lipid metabolism, agreeing with previous findings in the literature (Begriche et al., 2011). We found that fatty acid utilization for energy decreased, but that metabolism of other lipids, such as glycerolipids and linoleate, was dependent on the drug being administered. The only subsystem active in both a sexually dimorphic and treatment-induced manner was xenobiotic metabolism, further indicating that this group of reactions should be the focus of further investigation.

Our individual models suggest the presence of a core and unique metabolism both within each sex and between the sexes. Between the sexes, this unique metabolism involved mainly transport reactions in males and purine metabolism in females. Neither of these subsystems were present in any sex-specific TIDEs. This result suggests that though linking genes to reactions provides greater context for comparison, there are still possible systemic differences in male- and female-sourced cells that can result in differential metabolism elsewhere. With the averaged sex-specific models, differences in the consumption of bile acids and salts are seen between the sexes. Bile acids have been previously shown to be sex-specific in their amount and composition (Phelps et al., 2019), agreeing with our findings. Estrogen and testosterone, prominent sex hormones, have been shown to be inhibitors of the active sodium cotransporter of bile salts (Wishart et al., 2018); however, female sex hormone concentration in the blood fluctuates while it remains relatively constant in males (Briggs & Briggs, 1972). Additionally, women who take estrogen-based oral contraceptives clear specific drugs more efficiently than men (Miners, Attwood, & Birkett, 1983). These findings indicate that sex hormones may inhibit the reabsorption of drugs into the portal vein, decreasing concentrations in the liver. The deconjugating microbiome of the patients may also impact how bile acids and salts are recirculated, as previous work has shown that the gut microbiome is sex-specific (Kim, 2022).

There are important factors to note when considering the results of this analysis. First, we assume that gene expression levels are directly related to protein levels. There are other post-transcriptional and post-translational modifications that can alter protein abundance that are not available with the given data (Vogel & Marcotte, 2012). Though this assumption is not unique to this paper, it is nonetheless an important caveat when discussing its results. Additionally, our analysis here only considers one tissue type at a time; the cross-talk between organs and organ systems is a necessary consideration when evaluating toxicity as absorption, distribution, metabolism, and excretion by different organs can impact liver injury (Talevi & Bellera, 2022).

Sex as a biological variable continues to be a relevant consideration for any biological study. Many expression datasets do not have sex information recorded with the samples, decreasing our pool of potential data to be modeled as well as the quality of the data. An understanding of the necessity for sex as a variable will provide the foundation required to further the pursuit of precision medicine and drug therapy.

## Methods and Materials

### Adverse event reporting

We used AERSMine (Sarangdhar, M. et al, 2016) to evaluate the difference in adverse event reporting in the United States between the first quarter of 2004 to the third quarter of 2021. Only those reports with age and sex information were used, and patients between the ages of 15 to 65 were considered. Adverse events were considered “liver-related” if labelled with one of the following: “drug-induced liver injury”, “hepatotoxicity”, “hepatic enzyme increased”, “hepatic and hepatobiliary disorders nec”, “liver function analyses”, “hepatic and hepatobiliary disorders”, “hepatic failure and associated disorders”, and “hepatic enzymes and function abnormalities”. Once compiled, the percentage of total reports was calculated for each quarter and compared with a two-sided Mann-Whitney U test. Visualization was performed in R.

### Gene expression analysis

Differential expression between male and female samples from data sets were found in the Gene Expression Omnibus (Liver: GSE130991, Kidney: GSE36059, Brain: GSE5281). For those datasets without information on sex, male was assigned to those samples that were in the top 16% of the male-specific SRY gene and female to those in the bottom 16% (one standard deviation above and below the mean). Differentially expressed genes were then found using the limma package (Ritchie et al., 2015) and filtered to those genes that could be found in the Human1 metabolic network reconstruction. Those genes that were significantly different between male and female were assigned their log2 fold difference, and all other genes were assigned a log2 difference of 0.

### Biased metabolic subsystems

To find the presence of sex-biased metabolic subsystems, tasks inferred from differential expression (TIDEs) (Dougherty et al., 2021) and Human1 (Robinson et al., 2020) were used. TIDEs identifies all genes related to reactions in a user-defined “task” and assigns each reaction a weight based on the log2 fold difference in the expression of those genes. For genes with an “OR” gene protein rule, the highest fold difference will be used; for genes with an “AND” rule, the lowest will be used. Each fold difference is then averaged for a given task, and this average becomes its task score. This score is then compared to 1000 randomized task scores, calculated using randomly chosen log fold difference weights from other tasks. Significance was decided if p < 0.025 because it is a two-sided test. Tasks were defined as KEGG ortholog subsystems as assigned by the model Human1.

### Sex-specific models

Sex-specific liver models were created using Human1 (Robinson et al., 2020), gene expression data (GSE130991), and Reaction Inclusion by Parsimony and Transcript Distribution (RIPTiDe) (Jenior et al., 2020). Using gene expression data, RIPTiDe assigns linearly distributed weights between 0 and 1 to each reaction in Human1 with 0 being the reaction assigned the highest abundance gene. The sum of fluxes is then minimized as the objective function, and any reactions with 0 flux are pruned. The weights in the resulting model are reassigned inverse to the previous (1 referring to the reaction with the highest gene abundance) and the sum of fluxes is maximized. The model is sampled using these constrained flux distributions. This technique was performed with male and female liver data, with 804 (586 female/218 male) individual models and averaged models for male and female created and sampled.

## Data Availability

Code to reproduce this analysis is available at (https://github.com/ConnorMoore1/Sex-Spec-Hepato). Gene expression data available under GSE130991, GSE36059, and GSE5281.

## Author Contributions

CM and JP conceived the study. CM performed the computational modeling and data analysis. CM wrote the initial draft of the manuscript. CM, CH, and JP edited and wrote the final manuscript.

